# VirID: Beyond Virus Discovery - An Integrated Platform for Comprehensive RNA Virus Characterization

**DOI:** 10.1101/2024.07.05.602175

**Authors:** Ziyue Yang, Yongtao Shan, Xue Liu, Guowei Chen, Yuanfei Pan, Qinyu Gou, Jie Zou, Zilong Chang, Qiang Zeng, Chunhui Yang, Jianbin Kong, Yanni Sun, Shaochuan Li, Xu Zhang, Wei Chen Wu, Chunmei Li, Hong Peng, Edward C. Holmes, Deyin Guo, Mang Shi

## Abstract

RNA viruses exhibit vast phylogenetic diversity and can significantly impact public health and agriculture. However, current bioinformatics tools for viral discovery from metagenomic data frequently generate false positive virus results, overestimate viral diversity, and misclassify virus sequences. Additionally, current tools often fail to determine virus-host associations, which hampers investigation of the potential threat posed by a newly detected virus. To address these issues we developed VirID, a software tool specifically designed for the discovery and characterization of RNA viruses from metagenomic data. The basis of VirID is a comprehensive RNA-dependent RNA polymerase (RdRP) database to enhance a workflow that includes RNA virus discovery, phylogenetic analysis, and phylogeny-based virus characterization. Benchmark tests on a simulated data set demonstrated that VirID had high accuracy in profiling viruses and estimating viral richness. In evaluations with real-world samples, VirID was able to identity RNA viruses of all type, but also provided accurate estimations of viral genetic diversity and virus classification, as well as comprehensive insights into virus associations with humans, animals, and plants. VirID therefore offers a robust tool for virus discovery and serves as a valuable resource in basic virological studies, pathogen surveillance, and early warning systems for infectious disease outbreaks.

## Introduction

RNA viruses are renowned for their genetic and phenotypic diversity and ability to infect hosts ranging from animals, plants, fungi, to microbial organisms, sometimes with devastating health and economic consequences (Nicaise 2014). RNA viruses have caused human epidemics for millennia, with notable recent examples including human immunodeficiency virus (HIV; 1981) (Fauci 1988), SARS-CoV (2002) (Zhong, et al. 2003), pandemic H1N1 influenza (2009) (Smith, et al. 2009), MERS-CoV (2012) (Assiri, et al. 2013), Ebola virus (Western Africa, 2013) (Team 2014), Zika virus (2015) (Musso and Gubler 2016) and most recently SARS-CoV-2 (2019) (Zhu, et al. 2020). Importantly, approximately 70% of these pandemic-causing viral pathogens originate from wildlife animals or utilize arthropod vectors (Chan, et al. 2013), and many evaded surveillance systems before their emergence in human populations (Claas, et al. 1998; Ergönül 2006; Peiris, et al. 2007; Martina, et al. 2009). Additionally, RNA viruses pose significant threats to agriculture, particularly as epidemics in domestic animals and crops can jeopardize global food security (Mackenzie, et al. 2004; Untiveros, et al. 2007; Scholthof, et al. 2011; Lee 2015; Robilotti, et al. 2015; He and Krainer 2020). Understanding the diversity of RNA viruses and enhancing surveillance of those that pose threats to humans and economically important species are therefore endeavors of utmost importance (Carroll, et al. 2018; Lefrançois, et al. 2023).

Meta-transcriptomics (i.e., total RNA sequencing) provides a potentially unbiased survey of the genetic information from all types of organisms in biological samples and has transformed the detection and characterization of RNA viruses (Simmonds, et al. 2017; Greninger 2018; Shi, Lin, et al. 2018; Shi, Zhang, et al. 2018; Zhang, et al. 2019). This method provides efficiency and breadth in virus discovery compared to traditional cultivation techniques (Shi, et al. 2016; Chen, et al. 2022; Zayed, et al. 2022), which are often restricted by their reliance on cell culture growth (Huhtamo, et al. 2012; Shi, Lin, et al. 2018), and to PCR-based approaches that depend on prior knowledge of existing viral diversity (Culley, et al. 2003). As a consequence, meta-transcriptomics has become the primary tool for discovering RNA viruses (Greninger 2018), with the highly conserved RNA-dependent RNA polymerase (RdRP) that is essential for RNA virus replication serving a powerful universal genetic marker (Holmes 2009; Li, Shi, et al. 2015; Lam, et al. 2020; Edgar, et al. 2022; Mifsud, et al. 2022; Shi, et al. 2023).

As well as sequencing, the bioinformatics tools for detecting and characterizing virus sequences within metagenomic data sets have experienced major improvements. Initial workflow tools like ViromeScan (Rampelli, et al. 2016) and Taxonomer (Flygare, et al. 2016) detected viruses in samples by analyzing sequencing reads. Subsequent tools, such as Vipie (Lin, et al. 2017), ID-seq (Kalantar, et al. 2020), Lazypipe2 (Plyusnin, et al. 2023), and ViWrap (Zhou, et al. 2023), use contigs from *de novo* assemblies for virus identification, enhancing the length of query sequences and aiding the discovery of divergent viruses. With the advent of machine learning and deep learning, tools such as PhiSpy (Akhter, et al. 2012), VirSorter (Roux, et al. 2015), VirSorter2 (Guo, et al. 2021), and VIBRANT (Kieft, et al. 2020) were developed to differentiate viral from microbial sequences by analyzing gene and/or genomic sequences and features. Furthermore, methods like VirFinder (Ren, et al. 2017) and DeepVirFinder (Ren, et al. 2020) utilize the frequency of consecutive nucleotides (k-mers) in known viral and cellular genomes to identify DNA bacteriophage. More recently, specialized RNA virus discovery software such as VirBot (Chen, Tang, et al. 2023) have been developed. VirBot constructs databases of RNA virus protein families and employs profile Hidden Markov Models (pHMM) to identify distantly related viral sequences, exhibiting superior performance to other virus classification tools.

Despite these advancements, the bioinformatics tools for virus discovery continue to confront major challenges, such as a high false positive rate, the overestimation of viral diversity, and inaccurate virus classification (Dutilh, et al. 2021; Hegarty, et al. 2024). These issues are particularly problematic for RNA viruses due to their highly divergent genomes and distinct genomic structures which complicate identification using basic annotation methods (Drake 1993; Simon-Loriere and Holmes 2011). Additionally, current bioinformatics tools often lack the capability for the in-depth characterization of viruses, including the accurate identification of host associations, which impedes an understanding of their potential threat to public health and agricultural systems.

In response to these challenges, we developed VirID – a comprehensive and user-friendly software tool tailored for the discovery and characterization of RNA viruses from metagenomic data. VirID features a robust database of RdRP sequences and comprises three core modules: (i) RNA virus discovery, (ii) phylogenetic analysis, and (iii) phylogeny-based virus characterization. Benchmark tests on simulated data set revealed VirID’s high accuracy in profiling and classifying RNA viruses. In practical applications with real-world data sets, VirID demonstrated its capacity for virus discovery and for conducting thorough virome analyses.

## Materials and Methods

### RdRP Database

We established a database of 7080 representative RdRP protein sequences for RNA virus discovery and characterization. The RdRP sequences were derived from three independent sources, including a “backbone RdRP data set” (*N* = 5,384) (Shi, et al. 2016; Shi, Lin, et al. 2018), the NCBI RefSeq database (viral RdRP associated, *N* = 19,574), and the NCBI GenBank database (viral RdRP associated, *N* = 5,710,331). RdRPs from the RefSeq and GenBank databases were initially identified based on key annotation terms, including “RdRP”, “RNA-dependent RNA polymerase” and “polymerase” under the taxonomy “Riboviria”. Highly divergent RdRPs derived from environmental samples (Edgar, et al. 2022; Zayed, et al. 2022; Hou, et al. 2023) were excluded due to lack of confirmation. The RdRPs from NCBI were then compared to the backbone RdRP data set using Diamond v2.1.4 (Buchfink, et al. 2015), with an e-value threshold of 1e-5 and the ‘--ultra-sensitive’ parameter, and sequences with BLAST hit results were retained. The presence of key RdRP domains, specifically the highly conserved A, B and C sequence motifs, were examined using palmscan v1.0.i86linux64 (Babaian and Edgar 2022). The remaining sequences were clustered using CD-HIT v4.8.1 (Fu, et al. 2012) at 80% amino acid identity, with a representative then selected from each cluster. CD-HIT’s default parameters chose the longest sequences, which were then reviewed and corrected for any errors. This resulted in a final reference database of 7080 sequences that were then systematically organized into 24 RNA virus “superclades” based on broad phylogenetic relationships (Figure 1A)(Shi, et al. 2016; Shi, Lin, et al. 2018), and subsequently into 5 phyla, 20 classes, 28 orders, and 112 families based on both the phylogenetic relationships and ICTV taxonomy (Figure 1B, Supplementary Table S1).

**Figure 1.**
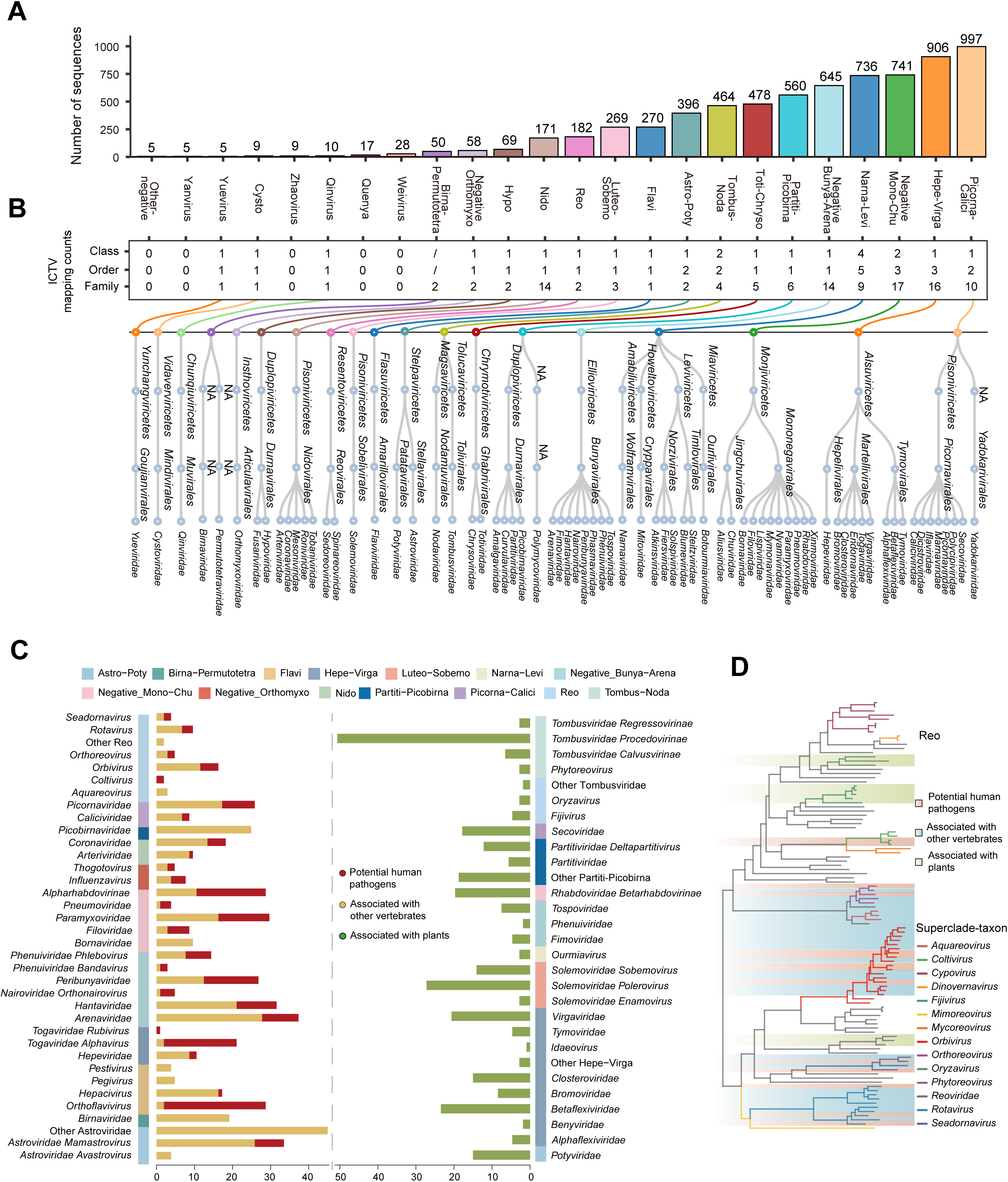
Overview of the RdRP reference database. (**A**) Compilation of representative RdRP sequences into 24 RNA virus superclades, categorized based on phylogenetic analysis. (**B**) Alignment of viral superclades with the ICTV taxonomic system, spanning virus class, order, and family levels. Numbers reflect the count of ICTV classifications mapped to each RdRP superclade, with specific ICTV representative mapping details provided. (**C**) The distribution of viral sequences across superclades and ICTV taxonomy, highlighting 551 vertebrate-related and 325 plant-related sequences. (**D**) Using the *Reoviridae* as a case study, the diagram presents a phylogenetic tree in which branch colors indicate taxonomy and background hues denote host association. Reference sequences are clearly annotated based on their positions within the phylogenetic tree.

For each of the RdRP sequences, information on host organism was initially retrieved from the Virus-Host Database (Mihara, et al. 2016) and confirmed by checking the original publications. Based on the host information and phylogenetic relationships, we categorized the RdRP sequences into four broad groups: (i) those infecting humans (including vector-borne viruses, *N* = 188), (ii) those infecting vertebrate animals (*N* =363), (ii) those infecting plants (*N* = 325), and (iv) all other host associations (Figure 1C). This information, together with virus phylogenetic relationships, was used to define specific host groups on phylogenetic trees (Figure 1D, Supplementary Figure S1) from which the host-association of the newly identified RdRP sequences could be inferred (See below).

### Processing and Assembly of Sequencing Reads

VirID performs an initial processing of the input read sequences. The quality control of reads was conducted using bbduk.sh (Bushnell 2014). To remove host and microbial ribosomal RNA (rRNA) sequences, the reads were then mapped, using Bowtie2 v 2.5.1 (Langmead and Salzberg 2012), against a reference rRNA database that contained a total of 505,405 rRNA non-repetitive sequences obtained from SILVA database v138.1 (Quast, et al. 2012) and the RDP-II database (Cole, et al. 2007). The remaining reads were assembled *de novo* into contigs using MEGAHIT v1.2.9 (Li, Liu, et al. 2015) and under default parameters. Only contigs longer than 600bp were retained for subsequent virus discovery and characterization.

### Discovery and Quality Control of RNA Virus Contigs

VirID discovers RNA viral contigs through a homology-based search approach. The assembled contigs were first compared against the RdRP reference database using the Diamond BLASTx program v2.1.4 with the e-value threshold of 1e−4 to identify potential RNA viral sequences. To remove false positives, potential viral contigs were subsequently compared against the NCBI Non-Redundant Protein Database (NR) with the e-value threshold of 1e−4 and the hits were annotated using TaxonKit v0.14.2 (Shen and Ren 2021). Contigs with only non-viral hits were removed, and the top hits of the remaining contigs were used to provide an initial taxonomic annotation for the viral contigs.

In addition to the standard quality control steps described above, VirID employs extra procedures to identify and remove false positives and erroneous sequences. To filter out potential endogenous virus elements (EVEs) with disrupted ORFs, we removed contigs whose RdRP-associated open reading frames (ORFs) were predicted to contain less than 200 amino acids by scanning the viral sequence in six possible reading frames. For ORF prediction, we included an option for 26 genetic code sets available in the NCBI database, with the standard code set as the default. To identify and control for misassembled contigs that contained both viral and non-viral sequences, the sequence was first compared using BLASTn (Camacho, et al. 2009) against a “non-viral” database, specifically a sub-set of NCBI Nucleotide Sequence Database (NT) that excludes viral sequences. Based on the position and length of hit, the target contigs were either partially corrected (i.e., in scenarios when “non-viral” regions appeared at either end of the contigs and were shorter than the “viral” region) or completely removed (all other cases).

### RNA Virus Species Identification and Quantification

For viral species identification, the RdRP-associated contigs were clustered using the all-to-all BLASTn method (Nayfach, et al. 2021), employing an 80% sequence identity and a 40% sequence length threshold to define approximate species-level taxonomic units. For each species-level group, the predicted RdRPs were compared against the NR database, and those with >90% identity and 70% coverage to existing RNA viral species were denoted “known” viral species, whereas those below these thresholds were considered as “new” viral species.

The sequence Reads Per Million (RPM) metric was used to evaluate the relative abundance of virus species within the sample. Clean reads are first mapped to all contigs associated with a virus species using Bowtie2 v2.5.1(Langmead and Salzberg 2012). The number of mapped reads is multiplied by 10^6^ and divided by the total number of reads to give the RPM value.

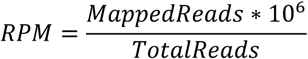

### Viral Sequence Alignment at the Superclade Level

For newly identified viral sequences, multiple alignment was performed before subsequent phylogenetic placement. Viral RdRP sequences were first classified into 24 superclades for sequence alignment and phylogenetic analysis, based on BLASTx comparisons (Camacho, et al. 2009) against the RdRP reference database. To accelerate the analysis of the Picorna-Calici superclade, we excluded contigs encoding proteins with fewer than 400 amino acids before alignment. This was necessary due to the Picorna-Calici superclade containing 533 reference sequences, which made each alignment iteration time-consuming. VirID utilizes the Amino Acid Consistency (AAC) index to quantify the similarity between two aligned amino acid sequences. AAC is calculated by dividing the number of identical amino acids (excluding gaps) by the total length of the aligned amino acid sequence.

To ensure the accuracy of multiple sequence alignments, we employed an iterative approach with Mafft v7.520 (Katoh and Standley 2013), followed by the removal of ambiguously aligned regions using trimAl v1.4 (Capella-Gutiérrez, et al. 2009). In each iteration, for each reference sequence (*R*^*i*^), its amino acid identity *r*^*i*^ is determined by the highest amino acid identity value compared to other reference sequences. For each potential viral sequence (*Q*^*i*^), its amino acid identity *q*^*i*^ is set as the maximum value when compared to all reference sequences. If *q*^*i*^ falls below the minimum value of *R* across the entire reference sequence set, the sequence *Q*^*i*^ is removed as a poor-quality sequence, and the alignment process is repeated. The pseudocode for eliminating false positives through multiple sequence alignment is provided in Supplementary Figure S2.

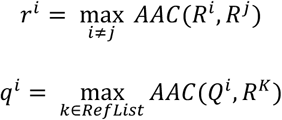

### Phylogenetic Placement

To optimize phylogenetic tree inference, we implemented an approach based on reference trees. We first estimated reference maximum likelihood trees in RaxML v7 (Stamatakis, et al. 2008), using the best fit amino acid substitution model (i.e., PROTGAMMALGF) selected by ProtTest (Abascal, et al. 2005). Subsequently, new sequences were integrated into these pre-built reference trees based on the alignment using the LG model in pplacer v1.1 (Matsen, et al. 2010), which accurately identifies their most likely topological positions. The output from pplacer, provided in JSON format, is then converted into the Newick tree format using guppy v1.1 (Matsen, et al. 2010), facilitating further phylogenetic analyses.

To evaluate the reliability of our phylogenetic placement, we performed leave-one-out cross-validation, focusing on tree recall, which measures the proportion of correctly recovered branches. For this analysis, we only included branches supported by a bootstrap confidence value exceeding 90%. We observed that the distribution of recall rates varied across superclades, likely influenced by the number of reference sequences and their similarity. In superclades containing more than 50 sequences, recall rates for most clades tended to concentrate around 0.75, with few deviations (Supplementary Figure S3). Additionally, in virus clades with a high density of sequences, such as Picorna-Calici and Partiti-Picobirna, this placement method maintained the fundamental stability of the tree structure, demonstrating robustness despite the addition of numerous new sequences.

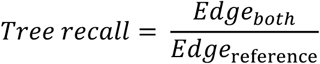

### Classification and Host Association Inference Based on Phylogenetic Analysis

The classification and host association of a query genome were inferred based on its placement within each RdRP superclade tree. We systematically annotated the evolutionary tree by traversing from the leaf nodes upwards, adhering to the principle that sibling nodes share the same labels. The labeling process for each leaf node followed the following criteria: (i) if the node corresponds to a known sequence, its label is assigned based on the existing annotation; (ii) if its sibling node is a known sequence, the leaf node inherits the label of its sibling; and (iii) if its sibling node is an internal node, we iteratively determined the label of that internal node before assigning it to the leaf node. For internal nodes, we conducted a systematic top-down traversal of all their daughter nodes. If all daughter nodes carry identical labels, the internal node adopts the same label as its descendants. However, if the daughter nodes have differing labels, the internal node is categorized under the lowest common taxonomic label for all those leaf nodes in this superclade. This structured approach ensures consistent and logical assignment of labels throughout each superclade tree.

Each potential viral sequence is assigned two types of labels: a taxonomic label that spans superclade-taxon to ICTV hierarchical classification and eventually down to the species level, and a host association label that categorizes the virus as related to humans, vertebrates, or plants. During the phylogenetic tree annotation, host association tags (represented as 0 or 1) are assigned to sequences. Specifically, if a viral sequence’s sibling node is identified as a human pathogen, the sequence is considered potentially relevant to humans only if the similarity exceeds 80%. For the ICTV classification within the phylogenetic tree, each internal node is labeled according to the lowest common taxonomic level of its descendant nodes. For sequences that remain unclassified, a default taxonomic hierarchy is assigned within the corresponding superclade. This classification system effectively categorizes potential viral sequences up to the genus level. For well-documented viruses, species-level classification is determined using the NCBI annotation file derived from comparisons within the NR database. This rigorous approach ensures accurate and detailed categorization of viral sequences across various levels of taxonomic and host associations.

### Benchmarking Preliminary Viral Contig Screening on Simulated Data Set

To evaluate the performance of VirID in the preliminary screening of RNA viral sequences, we constructed a simulated short-read shotgun metagenomic data set for benchmarking using CAMISIM (Fritz, et al. 2019). This data set included 15 samples from three categories: ‘known RNA viruses’, ‘new RNA viruses’, and ‘others’, with each category comprising 15 species. The ‘known RNA viruses’ and ‘others’ data sets were derived from 463 complete RNA virus genomes and 55 complete non-RNA virus genomes available in the NCBI RefSeq database, respectively (Supplementary Figure S4). The ‘new RNA viruses’ were subdivided into three levels based on their sequence similarity to the reference database and containing five species each: ‘new RNA virus L1’, ‘new RNA virus L2’, and ‘new RNA virus L3’, with similarity thresholds set at 0.70 ≤ similarity (L1) ≤ 0.90, 0.40 ≤ similarity (L2) < 0.70, and 0.20 ≤ similarity (L3) < 0.40, respectively. The sequences used to simulate ‘new RNA viruses’ were sourced from 299 high-quality RNA virus sequences obtained from recent publications (Feng, et al. 2022; Cui, et al. 2023; Hou, et al. 2023; Wang, et al. 2023), ensuring a comprehensive and challenging test environment.

The individual simulated data samples are provided in pair-end fastq format, with each file approximately 20 GB in size and each read 150 base pairs long. The distribution of the three data types within each sample is maintained at a consistent ratio of 1:1:1. Unlike other tools that require contigs, VirID accepts reads directly as input. To ensure consistent data standards across all tools, the contigs generated from VirID’s intermediate outputs are used as input for the other tools. Additionally, for uniformity in evaluation metrics, all tools have adopted VirID’s criterion of using contigs longer than 600bp.

We assessed the performance of three different RNA viral detection tools—VirSorter2, VirBot, and VirID—on a simulated data set using accuracy, precision, recall, and F1 score. These metrics focus on the number of correctly identified species in the tool’s output, rather than the number of sequences. The accuracy of each tool reflects the proportion of RNA viral species correctly identified from the total number of species. Recall measures the tool’s sensitivity by calculating the proportion of RNA viral species detected relative to the total number of RNA viral species in the data set. Precision indicates the specificity of the tool, defined as the proportion of RNA viral species detected out of the total number of species identified by the tool. The F1 score, the harmonic mean of precision and recall, provides a balanced measure of a tool’s overall effectiveness.

### Metagenomic Data Sets

To demonstrate VirID’s visualization capabilities, we analyzed 20 SRA libraries from a large scale animal study conducted in China (Cui, et al. 2023) that comprised 14 animal species across five provinces. In addition, we assessed the versatility of VirID through the analysis of 192 public libraries from seven studies, which included a wide range of samples from human swabs (Graf, et al. 2016) and diverse animal tissues (Chang, et al. 2020; He, Wang, et al. 2022; Shi, et al. 2022), as well as arthropod (Pettersson, et al. 2020), plant (Elmore, et al. 2022), and soil samples (Bender, et al. 2021). The public metagenomes utilized in this analysis were sourced from NCBI, with details provided in Supplementary Table S2, S3.

### Benchmark on Real Metagenomics Data Set

To evaluate the performance of VirID and VirBot for real-world data sets, we conducted a benchmark analysis using 20 wildlife sequencing libraries (Supplementary Table S2). VirBot utilized intermediate assembled contigs from VirID as input. For the contigs identified by VirID and VirBot, ORFs were translated using standard genetic code and annotated based on aligning predicted amino acid sequences to hidden Markov models (HMMs) from the Pfam-A database (https://pfam-legacy.xfam.org/) using HMMER’s hmmscan (http://hmmer.org/), with a minimum score threshold of 25. ORFs without hits in Pfam-A were further annotated using BLASTp against the NR protein database with an e-value threshold of 1e-4. We then categorized the top hit annotations of ORFs into three groups: (i) those contained RdRP protein, and those did not contain RdRP, but contained (ii) non-structural proteins other than RdRP, and (iii) structural proteins. To further compare the taxonomic classification performance of VirID and VirBot, we compared the classifications of contigs identified by VirID. We calculated the percentage of sequences classified at each of the seven taxonomic levels, from realm to genus. We benchmarked the runtime of VirID, VirBot, and VirSorter2 using 5 real-world sequencing samples with sequencing data size ranging from 2.6GB to 15.8GB (see Supplementary Table S4 for details). All runtime estimations were conducted in an environment running Ubuntu 22.04.2 LTS (GNU/Linux 5.15.0-116-generic x86_64) with four AMD EPYC 7643 48-core processors and 1.0 TB of memory. The programs were restricted to 32 threads, with no memory limitations. VirID used sequencing reads in ‘.fastq.gz’ format as input, while VirBot and VirSorter2 used intermediate assembled contigs from VirID, comprising sequences longer than 300 bp. To ensure a fair comparison, we used contigs as the starting material when performing comparisons among the three methods. Additionally, we estimated the computational runtime for the unique steps in the VirID analysis pipeline, including (i) read assembly and (ii) phylogenetic inference, recognizing that these steps could not be directly compared with the other methods due to the absence of equivalent processes.

### Parameters and Visualizations

VirID offers three user-selectable modes: ‘end to end’, ‘assembly and basic annotation’, and ‘phylogenetic analysis’. The ‘end to end’ mode executes both ‘assembly and basic annotation’ and ‘phylogenetic analysis’ sequentially. The ‘assembly and basic annotation’ mode processes sequencing data up to the point of producing annotated contigs, but it does not include phylogenetic analyses. Conversely, the ‘phylogenetic analysis’ mode begins with contigs and carries out subsequent processing from there.

During the ‘assembly and basic annotation’ stage, users can opt to remove mis-assembled portion of a virus contig, a process that may require longer time and significant memory resources. Additionally, an ultrasensitive mode is available for analyzing RdRP libraries that enhances the detection of a broader range of potential viral sequences. In the ‘phylogenetic analysis’ phase, users can choose to eliminate redundant sequences and customize the amino acid length thresholds for specific superclade of interest. VirID employs several tools for visualizations: sankey diagrams, stacked histograms, and phylogenetic tree diagrams are generated using the R packages networkD3 (Allaire, et al. 2017), ggtree (Yu, et al. 2017), and ggplot2 (Wickham 2011), respectively. Further visualizations are created using Matplotlib v3.7.4 (Hunter 2007) and Seaborn v0.13.0 (Waskom 2021), enabling comprehensive graphical representations of the data.

## Results

### VirID Workflow

VirID is a user-friendly command-line tool designed for the detection and detailed characterization of RNA viruses from metagenomic data. The workflow is divided into three stages: RNA virus discovery, phylogenetic analysis, and phylogeny-based virus characterization (Figure 2). In the first stage, putative RNA virus genomes containing the RdRP gene are identified through a homology approach, checked for false positives and contamination, and clustered based on sequence similarity at the species level. This process yields high-quality genomic sequences and related information such as sequence length, closest relatives, and abundance levels. In the second stage, VirID conducts high-quality sequence alignment and comprehensive evolutionary analysis across the full diversity of the RNA virosphere: this further removes low-quality sequences and reveals the phylogenetic position of all newly discovered viruses. In the final stage, based on these evolutionary analyses, VirID provides precise classification information and predicts potential viral associations with humans, vertebrates, and plants.

**Figure 2.**
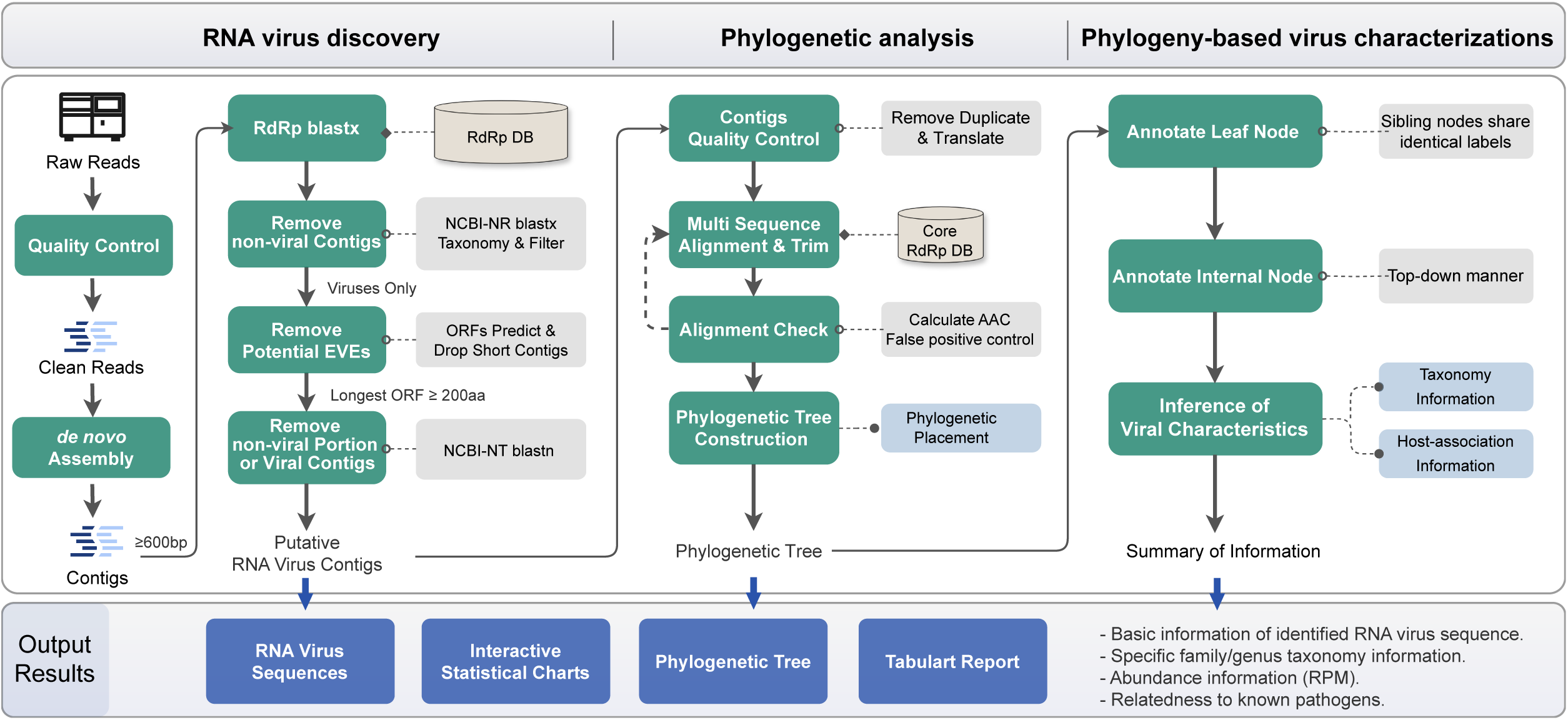
VirID framework for identifying RNA viruses from sequencing samples. The VirID framework for automated RNA virus detection, which comprises three main stages: (i) RNA virus discovery, (ii) phylogenetic analysis, and (iii) phylogeny-based virus characterization. It produces outputs that include viral sequences, phylogenetic trees, and comprehensive information including sequence length, best match of BLASTx comparison, virus classification, and host association.

### Analysis Results Output in VirID

VirID conducts thorough data analyses and offers robust visualization tools, enabling an intuitive and comprehensive presentation of results. VirID organizes its output into three main folders: (i) ‘assembly and basic annotation’ for intermediate files related to RNA virus discovery, (ii) ‘phylogenetic analysis’ containing well-labelled trees for various superclades, and (iii) ‘results’ for other figures and tables. It generates tables, fasta format sequences files, web files, and figure files (pdf format) to suit various analytical needs. We demonstrated these features here using a data set from 20 wildlife sequencing libraries that includes bats, insectivores, pangolins, pika and rodent samples (Cui, et al. 2023) (Figure 3, Supplementary Table S2). In the RNA virus discovery segment, VirID estimates Reads Per Million (RPM) values for each potential viral species to assess their relative abundance. These values are displayed in a sankey plot on an HTML webpage, illustrating RPM distributions across RdRP superclades and NCBI taxonomy at multiple taxonomy levels—phylum, class, order, family, genus, and species (Figure 3A). VirID also categorizes potential viral sequences into one or two of four host-association groups: human, vertebrate, plants, and others (Figure 3B). This information is provided in color-coded trees that contain information on both phylogenetic relationships and potential host associations (Figure 3C). Additionally, VirID outputs high-quality viral genome sequences in fasta format (Figure 3D) and all other relevant information, including sequence length, highest BLASTx match, virus classification, association with principal hosts, in multidimensional tables (Figure 3E, Supplementary Table S5, S6).

**Figure 3.**
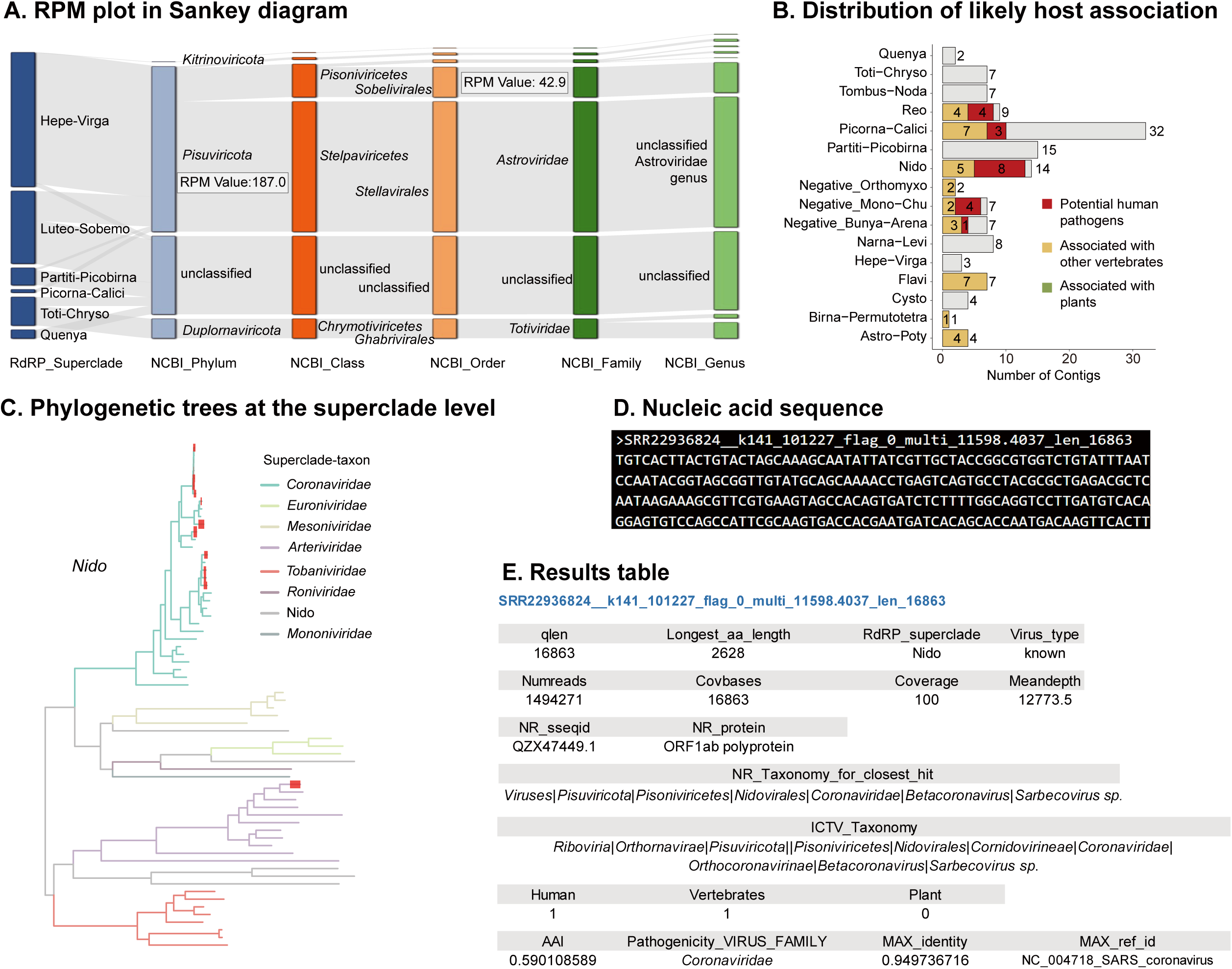
Visualization of outputs from VirID. (**A**) Distribution of Reads Per Million (RPM) of potential viral sequences from SRA sample SRR22936818, categorized by RdRP superclades and corresponding NCBI taxonomic ranks. (**B**) The host associations of all identified virus sequences based on their placement in the phylogenetic tree. (**C**) A phylogenetic tree is used to distinguish different viral lineages and colour-coded by taxonomic group. In this case, newly identified virus sequences in the Nido superclade are highlighted in red at the end of the branches, emphasizing their significance. (**D**) Complete nucleic acid sequences for detailed analysis and verification. (**E**) Detailed information for each identified viral sequence, including host association, genomic details, and taxonomic classification.

In the example of the 20 wildlife sequencing libraries VirID identified 129 potential RNA viral contigs belonging to 107 various species, including 87 new virus species, spanning 26 families and 15 superclades (Supplementary Figure S5A). Overall, 54 of these species can be assigned at the family level, with the remainder falling outside known families. Eleven of these species are potentially relevant to humans, including Wufeng Niviventer niviventer orthohantavirus 1, Severe acute respiratory syndrome-related coronavirus, Rotavirus A, and Pangolin respirovirus (Supplementary Figure S5B, Supplementary Figure S6). Additionally, several viral species associated with vertebrate infections were identified, including those within the genera *Alphacoronavirus*, *Alphainfluenzavirus*, and *Mammarenavirus*. The remaining 88 viral species are most likely derived from diet, parasites, and other microbial cellular organisms within the principal host, because they were identified as plant viruses (e.g. members of the *Tombunsviridae*) or closely related to arthropod (e.g. *Iflaviridae*) or fungal viruses (e.g. *Mitoviridae*).

### Benchmarking VirID using Simulated Data

We initially evaluated VirID alongside two other tools—VirSorter2, VirBot—using a collection of simulated samples (N = 15) that included a diverse mix of RNA viruses and other species. These read data samples were generated using the CAMISIM simulator, drawing on source genomes and sequences from both the NCBI RefSeq database and recent studies. The RNA viruses in the CAMISIM pool were comprised of four categories: ‘known RNA viruses’, ‘new RNA virus L1’, ‘new RNA virus L2’, and ‘new RNA virus L3’, representing different similarity levels to the reference database sequences.

Three tools were compared across 15 data samples, including the average accuracy, precision, recall, and F1 score metrics (Figure 4A). VirID achieved the highest F1 score of 0.9334, with a standard error of ±0.0180. Its recall, at 0.8822 with a standard error of ±0.0301, was close to VirBot (0.8822 ± 0.0247). Notably, VirID’s precision was perfect across all 15 samples, with an absence of false positives. The three tools were further evaluated for their performance on ‘new RNA viruses’ that shared less than 90% amino acid identity with those described in NCBI database (Figure 4B) and ‘known RNA viruses’. VirID continued to perform well in identifying ‘new RNA viruses’, again with zero false positives (Supplementary Figure S7).

**Figure 4.**
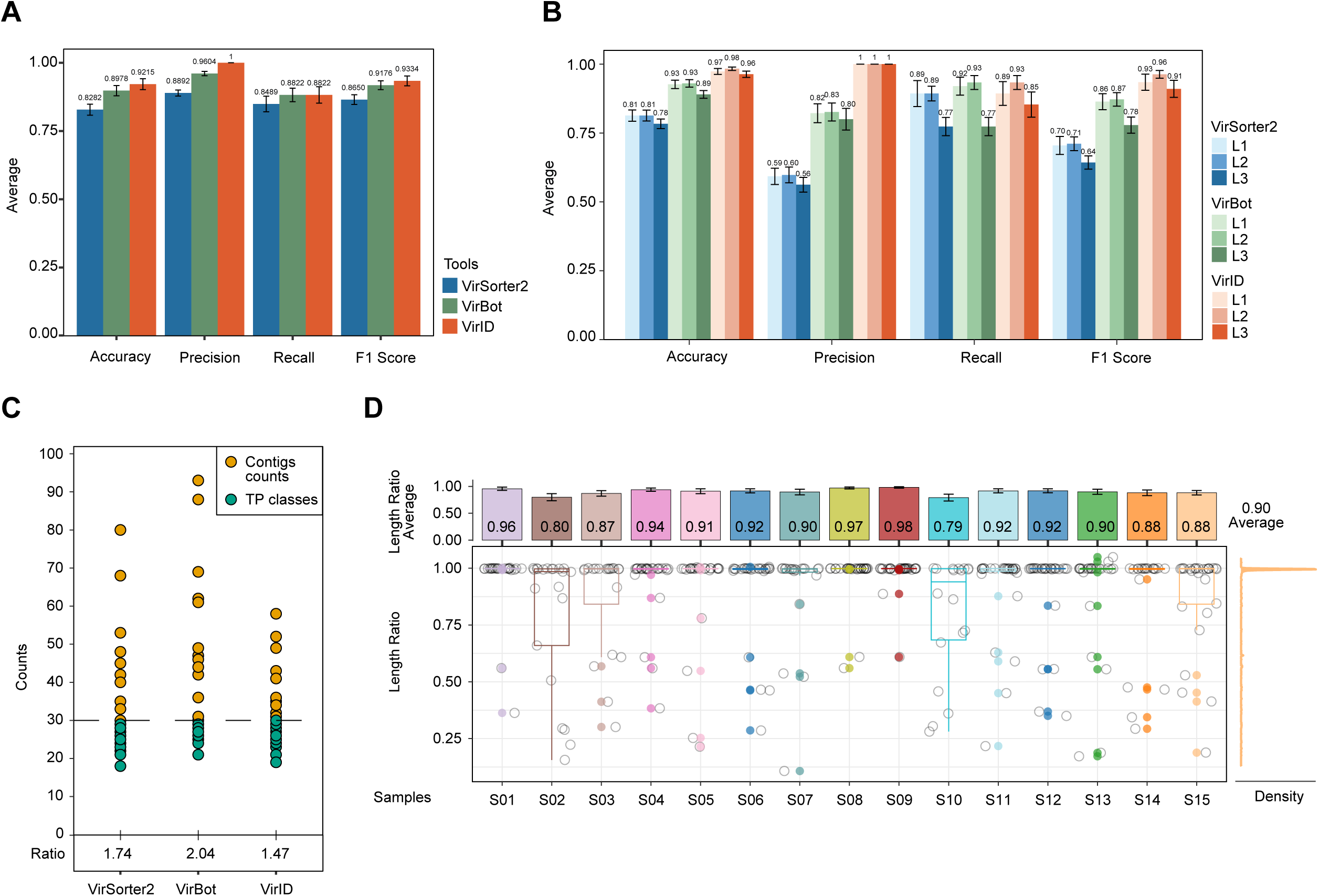
Benchmarking of three tools for virus discovery using simulated data. (**A**) Performance of three bioinformatics tools – VirID, VirBot, and VirSorter2 – across 15 simulated samples, evaluating average accuracy, precision, recall, and F1 score. Error bars represent standard errors. Notably, the training data sets for VirSorter2 and VirBot are primarily based on all viral genomes in the NCBI RefSeq database and a curated collection of viral proteomes, respectively. (**B**) The three bioinformatics tools were further assessed for their specific ability to identify ‘new RNA viruses’ at various levels of classification. (**C**) Effectiveness of each tool in identifying RNA virus sequences across 15 simulated samples. Yellow dots represent the number of RNA virus sequences identified, while green dots indicate the number of corresponding species-level taxa. (**D**) Ratio of the lengths of sequences identified by VirID in various simulated data samples compared to the length of the corresponding gold standard sequence. Only sequences that uniquely correspond to gold standard sequences are considered.

We also analyzed whether the number of viral sequences or species revealed by each tool matched that of expected species count in the simulated data (Figure 4C). VirID’s estimation of contigs (39, average) and species (27, average) were the most closely aligned with the 30 true species samples, followed by VirSorter2 (average of 45 contigs and 26 species) and then VirBot (53 contigs and 27 species). VirID achieved a ratio of 1.47 for the number of estimated sequences relative to the number of true species. This ratio was the lowest among all tools, indicating that VirID’s output most accurately reflect the actual number of virus species.

The integrity of the RNA viral sequences revealed and processed by VirID was assessed using the CAMISIM simulator’s ‘gold standard’ contigs as a benchmark. These standard contigs represent idealized assembly results of the simulated reads, providing a baseline for evaluating the detected virus sequences. We aligned the putative viral contigs with these ‘gold standard’ contigs, selecting the best matches based on coverage and identity. The average completeness for contigs in 10 samples exceeded 90%, with an overall average completeness of 90.80% across all 15 samples (Figure 4D). This meets CheckV’s criterion for ‘high quality completeness’ (Nayfach, et al. 2021).

### Benchmarking VirID on Real-world Sequencing Libraries

We conducted a benchmark analysis of the VirID and VirBot RNA discovery tools using the 20 wildlife sequencing libraries described above (Supplementary Figure S8). VirID and VirBot both identified 212 contigs (Supplementary Figure S8A). VirBot detected an additional 438 unique contigs, of which 240 were short contigs (<200 amino acids) and the remainder primarily encoded non-RdRp proteins. In contrast, VirID identified 23 additional contigs, all of which encoded RdRPs (Supplementary Figure S8B). This suggests that VirBot identifies more viral contigs overall, while VirID gives a more accurate representation of viral richness. Additionally, we compared the percentage of annotated contigs from the final 129 contigs obtained by VirID. Since VirID uses phylogenetic analysis for classification, it achieved more detailed viral classification across various taxonomic levels (Supplementary Figure S8C).

In regard to computational runtime estimation based on 32 threads of the AMD EPYC 7643 CPU, VirID (median 15.7 minutes, 11.3 - 28.4 minutes) and VirBot (median 5.2 minutes, 1 - 14.1 minutes) demonstrated similar performance, while VirSorter2 (median 141.9 minutes, 3.4 - 216.8 minutes) was significantly slower (Supplementary Figure S8E). For the unique steps in VirID, read assembly took a median of 92.5 minutes (4.5 – 126.2 minutes), with runtime positively correlated with the sequencing depth of the sample (Supplementary Figure S8D). Phylogenetic inference had a median runtime of 1.3 hours (0.3 – 16.9 hours), with the longest time spent on sequence alignment (Supplementary Figure S8F). The longest runtimes (16.3 and 16.9 hours) were observed in data sets containing members of the Picorna-Calici clade, which included a total of 533 reference viral sequences. The CPU runtime for phylogenetic inference could be significantly reduced by improving the speed of sequence alignment.

### A Re-Analyses of Previously Published Data Sets

We next assessed the performance of VirID by analyzing 192 meta-transcriptomic SRA libraries from seven distinct virome studies, covering a range of sample types including clinical (human), wild birds, Malayan pangolins, Qinghai voles, seabird ticks (*Ixodes uriae*), soybean fields, and peat soil samples (Supplementary Table S3). VirID significantly improved viral discovery. A total of 1,283 RNA viral species were identified in these data sets, with the soil samples displaying the highest average virus species richness at 88.5 species per sample, followed by Qinghai vole at 2, and then seabird tick at 1.9. Importantly, 1,202 novel RNA viral species were identified. Among these, the highest number of novel viruses (*N = 740*) was found in the soil samples, with 56% of these viruses categorized within the Narna-Levi superclade. At the family level, the *Picobirnaviridae* contained the most viral species (*N = 164*), followed by the *Tombusviridae* (*N = 83*) and *Mitoviridae* (*N = 69*), although many virus species (54.5%) could not be assigned at family level (Figure 5A).

**Figure 5.**
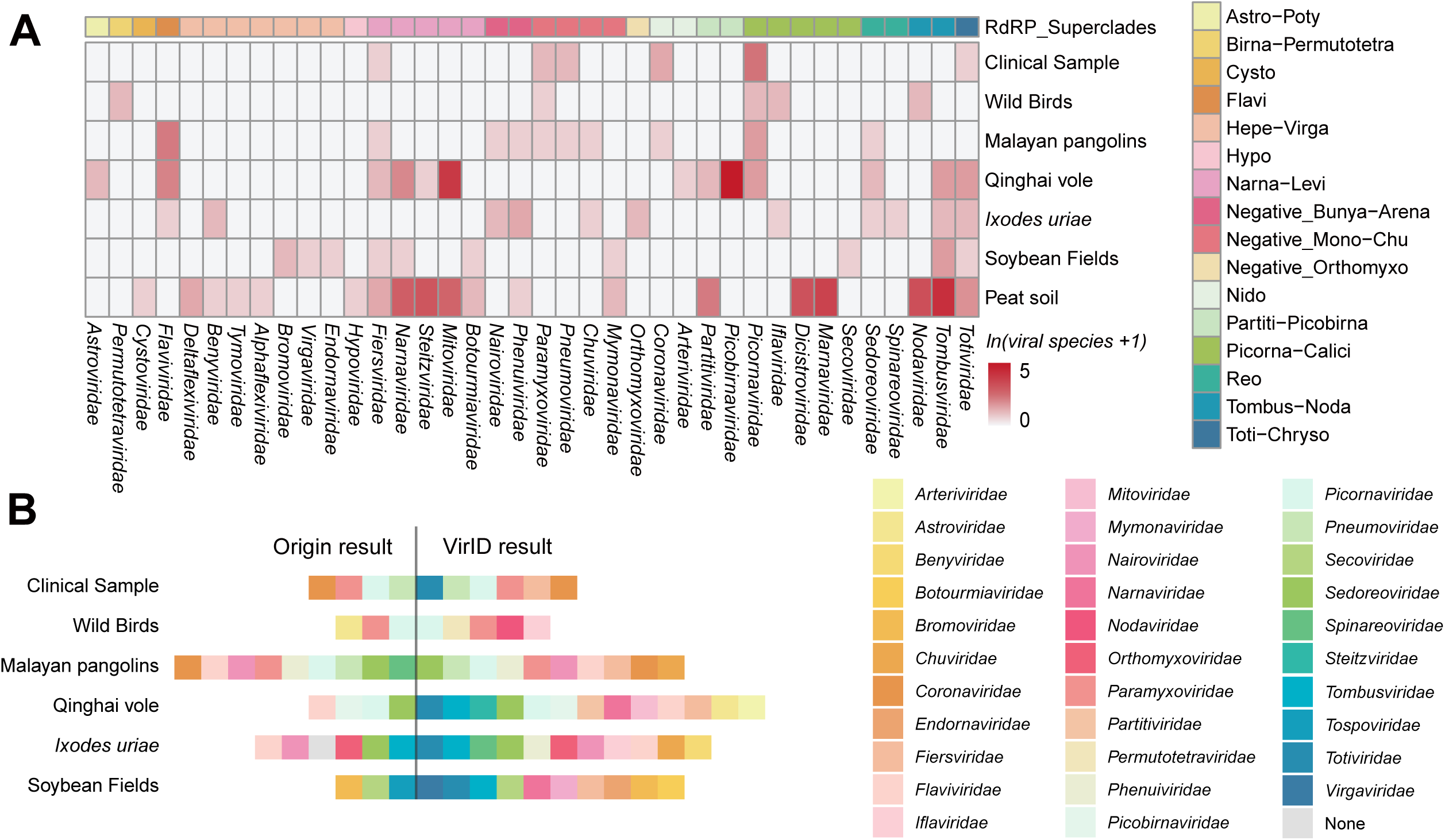
Reclassification of viruses using VirID on real-world data. (**A**) Distribution of virus species across different superclades and families identified by VirID in previously published data sets, including human swabs (Graf, et al. 2016), wild birds (Chang, et al. 2020), Malayan pangolins (Shi, et al. 2022), Qinghai voles (He, Wang, et al. 2022), seabird ticks (Pettersson, et al. 2020), soybean fields (Elmore, et al. 2022), and soil samples (Bender, et al. 2021). (**B**) The utility of VirID across a broader spectrum of RNA virus families. Virus families documented in the original publications are displayed on the left, while those identified in this study are shown on the right.

We next evaluated the number of families identified and compared these with previous studies, omitting any with incomplete classification information. Our analysis revealed an average increase of five viral families, with notable differences particularly observed in the vole and soybean data sets (Figure 5B). In addition, VirID provided a more precise classification system for the target viruses. For instance, viruses that were previously classified only at the kingdom level, such as Bulatov virus, Fennes virus, Ronne virus, and Vovk virus from the seabird tick data set, were now classified at the genus level (Supplementary Table S7).

VirID also provides information on whether the viruses observed are associated with vertebrates, plants, or have the potential to infect humans. For example, in the Qinghai vole data set we identified 180 species that were likely associated with voles, including *Mamastrovirus* (N = 1), *Hepacivirus* (*N = 5*), *Pegivirus* (*N = 2*), *Arteriviridae* (*N = 1*), and *Picobirnaviridae* (*N =164*), although the host association for the *Picobirnaviridae* remains uncertain (Sadiq, et al. 2024). Additionally, 250 virus species were likely associated with food sources, parasites, and symbionts within these hosts, including Narna-Levi (*N = 163*), Partiti-Picobirna (*N = 74*) and Toti-Chryso (*N = 6*) (Figure 6A). Overall, vertebrate-associated viruses accounted for a median of 66% of the total non-ribosomal RNA in host samples, ranging from 30.2-73.6%, while other viruses accounted for 23.3%, with a range of 10.5-67.9% (Figure 6B). Notably, we identified two species, Hepatovirus D and Rotavirus A, that are potential human pathogens. Hepatovirus D was described in the original publication (He, Wang, et al. 2022), while Rotavirus A was newly identified here. These viruses were discovered because they cluster with known human pathogens at over 80% sequence identity with these viral contigs.

**Figure 6.**
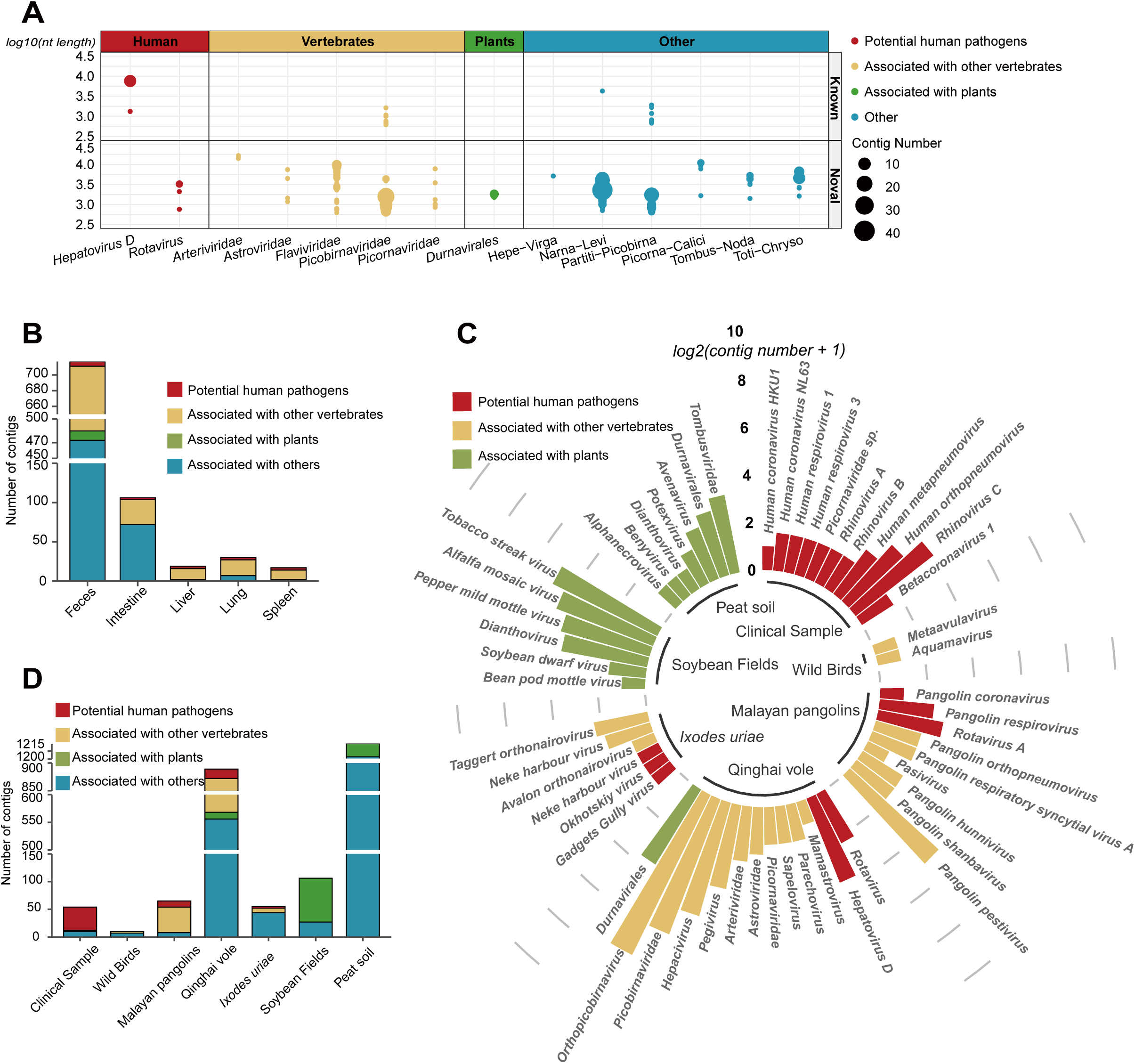
Virus-Host associations revealed by VirID. (**A**) Number of known and unknown virus sequences related to viruses of humans, vertebrates, plants, and other hosts as identified by VirID in the Qinghai vole case study. (**B**) Count of virus sequences associated with different hosts in various tissue and fecal samples from voles identified by VirID. (**C**) Number and taxonomy of virus sequences related to different hosts in various real-world samples identified by VirID. (**D**) Number of virus sequences related to viruses of different hosts in a variety of real-world samples identified by VirID.

Similarly, we identified 17 potential human pathogens, primarily within the clinical sample data set. This included potential human pathogens such as Pangolin coronavirus HKU4, Rotavirus A, and Pangolin Respirovirus identified in pangolins, as well as the vector-borne Gadgets Gully virus, Okhotskiy virus, and Neke harbour virus from the seabird tick samples. This is consistent with previous publications. Additionally, plant-infecting viruses were identified in the vole (*N = 5*, likely diet-related), soybean (*N = 6*), and soil (*N = 24*) samples (Figure 6C, Supplementary Table S8). In the Soybean field data set, we detected 79 contigs from six viral species associated with plants, constituting about 74.5% of the total non-ribosomal RNA, including tobacco streak virus, soybean dwarf virus, pepper mild mottle virus, bean pod mottle virus, and alfalfa mosaic virus (Figure 6D).

## Discussion

Herein, we present VirID, an RNA virus discovery platform specifically designed for Linux servers. VirID integrates phylogenetic analysis into the identification and characterization of RNA viruses, thereby substantially improving virus discovery. The platform employs an iterative scoring strategy in its alignment processes to minimize false positives and incorporates a phylogenetic placement algorithm for the rapid and stable phylogenetic tree inference. As a result, users can accurately estimate RNA virus diversity, classify each identified virus with high precision, at the same time obtaining insights into host associations which enable the potential threat to human, animal and plant health to be evaluated. The wealth of information generated by this platform will be invaluable for researchers involved in early warning of infectious diseases across diverse settings.

Our results show that VirID provides the most precise estimation of viral species composition, while other programs tend to overestimate genetic diversity. This overestimation may occur because some methods consider all virus-associated contigs, including those with low coverage and fragmented genomes, potentially counting them as multiple species. To depict diversity more accurately, it is essential to focus on regions of the genome shared by the majority of viruses, thereby minimizing overestimation (Beerenwinkel, et al. 2012; García-López, et al. 2015). The RdRP gene is an excellent candidate for several reasons. First, all RNA viruses, with the exception of RNA satellite viruses, contain the RdRP (Shi, et al. 2016; Shi, Lin, et al. 2018; Zayed, et al. 2022). Second, the RdRP protein is the most conserved gene in the RNA virus genome, and often comparable across different virus classes or even phyla (Venkataraman, et al. 2018; Mönttinen, et al. 2021). Therefore, focusing solely on RdRP contigs is key to accurately determining viral diversity within a sample.

Compared to other virus discovery programs, the phylogenetic analysis feature of VirID provides a central analytical component. In particular, it provides an automated and reliable method for accurately classifying previously undescribed RNA viruses, especially those that are highly divergent in sequence. Traditionally, the approach to classifying new viruses involved BLAST analyses and assigning taxonomic position based on the closest related hits (Lin, et al. 2017; Zhao, et al. 2017; Plyusnin, et al. 2023). However, this method may counter the guidelines laid down by the International Committee on Taxonomy of Viruses (ICTV) who apply varying criteria for virus taxonomy at the species and genus levels (King, et al. 2012; Simmonds, et al. 2017; Lefkowitz, et al. 2018). Additionally, the lack of a clear definition of what levels of genetic similarity differentiate higher taxonomic ranks complicates the assignment of viruses to new families or orders when protein identity is below 40%. More importantly, some reference sequences are incorrectly classified, which can introduce errors into the classification process. For instance, a sequence from soil metagenomic data (MN035928), from a divergent member within the order *Bunyavirales*, was mistakenly labeled as belonging to the genus *Arenavirus* within the *Arenaviridae* (Starr, et al. 2019). As a consequence, subsequent discoveries of similar viruses (OQ715420)(Chen, Hu, et al. 2023) were also mislabeled as *Arenaviridae*. Such misclassifications could be avoided with robust and reliable phylogenetic analyses.

Phylogenetic analysis is also central to the inference of host associations. Viruses from similar host categories tend to cluster together, forming what is known as a phylogenetic monophyly, indicative of host structure in virus phylogeny (Kitchen, et al. 2011; Shi, et al. 2016; Shi, Lin, et al. 2018; French, et al. 2023). This pattern holds true across different types of viruses, such as vertebrate-specific viruses, arthropod-borne viruses, and plant viruses, in which host-associated phylogenetic monophyletic groups are identifiable at various taxonomic levels—ranging from the family level (e.g., *Picornaviridae*) to the genus level (e.g., *Alphavirus*), and even within specific genetic lineages in a single genus (e.g., mosquito-borne and tick-borne virus groups within genus *Flavivirus*). Due to the varying scales and complexity of these host-associated groups, phylogenetic analysis serves as a valuable tool for exploring these relationships (Shi, et al. 2022; Cui, et al. 2023; Wang, et al. 2023). However, assigning viruses to other host groups such as arthropods, nematodes, and even basal eukaryotes (such as parasites) remains challenging due to the scarcity of relevant virus data on these hosts. In addition, a limitation to all virus discovery tools, including VirID, is that they may not identify vertebrate-associated viruses if they occupy phylogenetic positions not previously linked to vertebrate hosts.

The ability to infer host associations is highly relevant in disease monitoring programs in which emerging pathogens are continuously identified and evaluated (Carroll, et al. 2018; Carlson, et al. 2022; Ko, et al. 2022). Numerous viruses are discovered in sequencing efforts, but not all are relevant to the infection of principal host or impact health. Indeed, those relevant to disease often constitute only a small fraction (Liang and Bushman 2021). For example, in a survey of over 1941 game animals across China, more than 1000 viruses were identified, yet only 102 were linked to mammalian infections, and even fewer (N = 21) were considered to pose a significant risk of infecting humans or other animal species (He, Hou, et al. 2022). Similarly, a recent meta-transcriptomics study of 2438 mosquitoes in China revealed that among the 564 RNA virus species detected, 393 were likely associated with mosquitoes, but only 7 were linked to mammalian infections (i.e., arboviruses) (Pan, et al. 2024). Thus, discerning potential host associations is crucial for assessing the public health or economic impact of discovered viruses (Rahman, et al. 2020; Bernstein, et al. 2022; Lefrançois, et al. 2023).

Our study is subject to several limitations. First, while the phylogenetic placement method has accelerated the estimation of evolutionary trees, the self-iterative multi-sequence alignment process remains time-consuming. Second, VirID primarily relies on the RdRP to denote homology and is therefore unable to identify sequences that do not contain RdRP sequences or that are too divergent to be detected in homology-based searches (Telesnitsky and Goff 2011; Hu and Hughes 2012). Consequently, for segmented viruses, the full genome assembly may be incomplete. Third, VirID does not provide strain or genotype level typing which often depend on more variable parts of viral genomes (Yang, et al. 2020; Liao, et al. 2022). Finally, the RdRP database is continuously expanding, particularly with the addition of data from environmental samples (Wolf, et al. 2020; Chen, et al. 2022; Edgar, et al. 2022; Zayed, et al. 2022; Hou, et al. 2023). However, since these environmental RdRP data are unverified, and this study focuses on human, animal, and plant health, they are currently excluded from our database. Nevertheless, given the flexibility of our RdRP database structure, these data can be readily incorporated to facilitate future broader scale analyses.

## Data Availability

VirID is freely available as an open-source Python code at https://github.com/ZiyueYang01/VirID. And the newly identified virus sequences from this study are available under the link: https://github.com/ZiyueYang01/VirID/blob/main/data_res/Novel_RNA_viral.fasta

## Supplementary Data

Supplementary Data are available at MBE Online.

## Author Contributions

Conceptualization, EC Holmes, DY Guo and M Shi; Methodology, ZY Yang, YT Shan, X Liu, GW Chen, YF Pan, DY Guo and M Shi; Investigation, ZY Yang and YT Shan; Data Collection and Processing, QY Gou, J Zou, ZL Chang, Q Zeng, CH Yang, JB Kong, WC Wu, DY Guo and M Shi; Writing – Original Draft, ZY Yang and YT Shan; Writing – Review and Editing, All authors. Funding Acquisition, CM Li, H Peng, DY Guo and M Shi; Resources (computational), SC Li, X Zhang, DY Guo and M Shi; Supervision, CM Li, H Peng, DY Guo and M Shi.

