## Supplementary Figures for "VirID: Beyond Virus Discovery - An Integrated Platform for Comprehensive RNA Virus Characterization"

**Content**

Supplementary Figures S1-S8

Supplementary Tables S1-S8

### Supplementary Figures

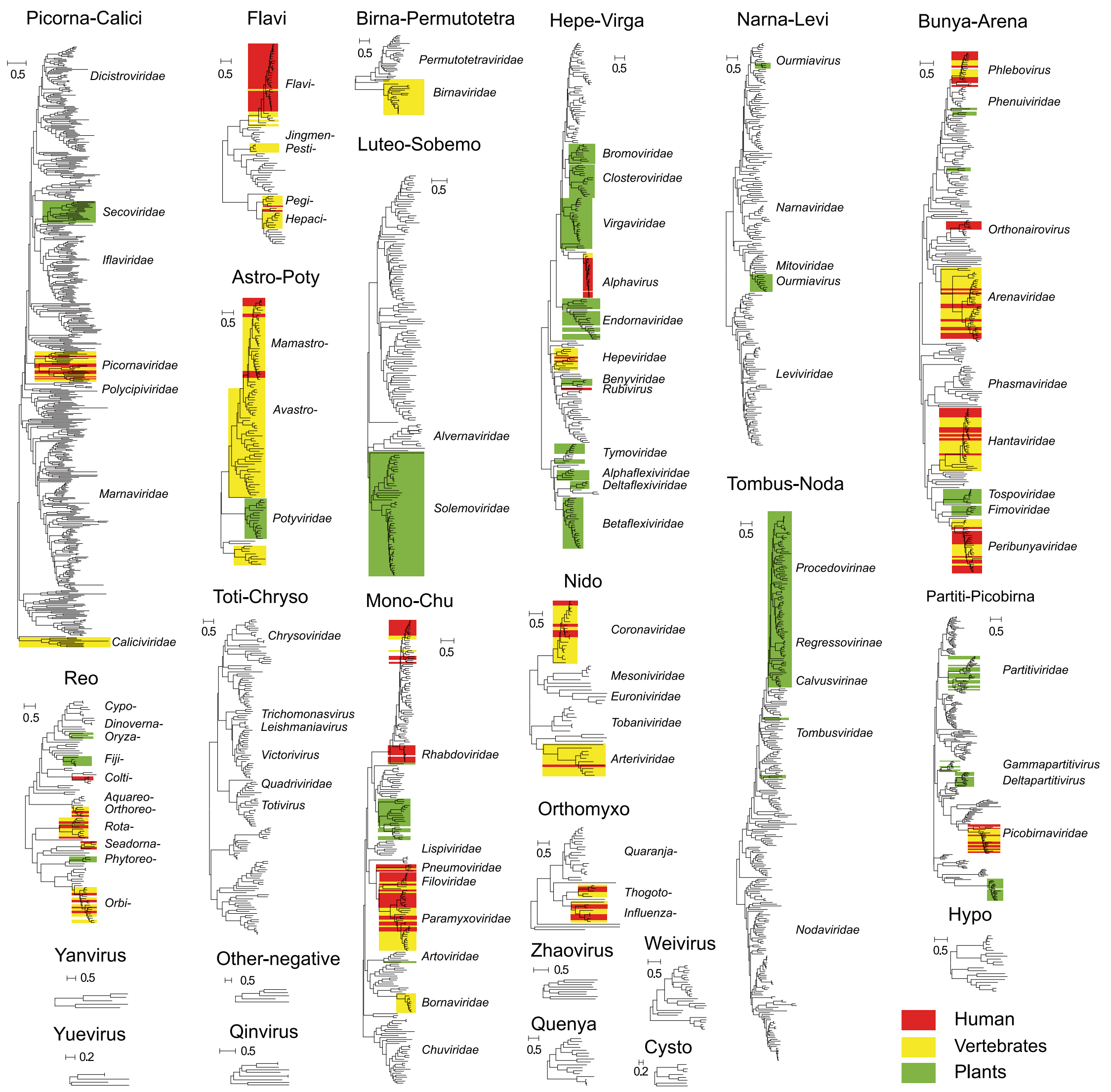


**Supplementary Figure S1. Host associations of reference viruses in 24 superclades.** A total of 24 phylogenetic trees where background shading signifies host type (i.e., human, vertebrates, plants). Reference sequences are annotated according to their phylogenetic positions within each phylogeny.

```
Algorithm 1:  Delete false-positive sequences
input  :  Ref seqs refList , query seqs queryList and Alignment res
          Alignment
output:   False-positive seqs delList
Funcion  AAC(strI,strJ) :
  | For num in (0,len(strI)) and strI != strJ ;
  | if strI[num] == strJ[num] and strJ[num] != gap: count+=1;
  | return count/len(strI);
RefAACList ← element e of refList : max {AAC( e,refList )};
minRefAAC ← min RefAACList;
for queryI in queryList do
  | maxQuery ← max {AAC( queryI ,refList )} ;
  | if maxQuery < minRefAAC then
  |   | delList ← delList + queryI
  | end
end
return delList
```

**Supplementary Figure S2. Pseudocode for eliminating false positives.** The AAC function is used to calculate the consistency between two amino acid sequences. After each multiple sequence comparison, VirID calculates the consistency between query sequence and the reference sequence to obtain false positive sequences.

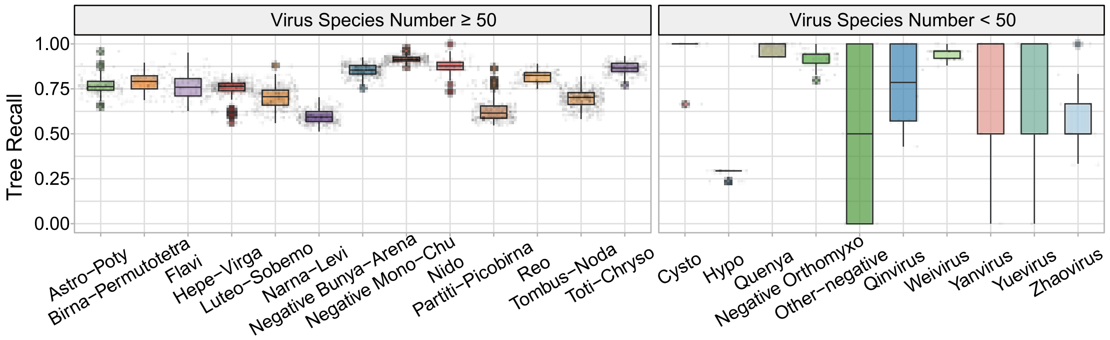


**Supplementary Figure S3. Accuracy of the VirID placement method for different superclades.** The performance of superclades performing leave-one-out cross-validation, evaluating tree recall. Each point represents one experiment.

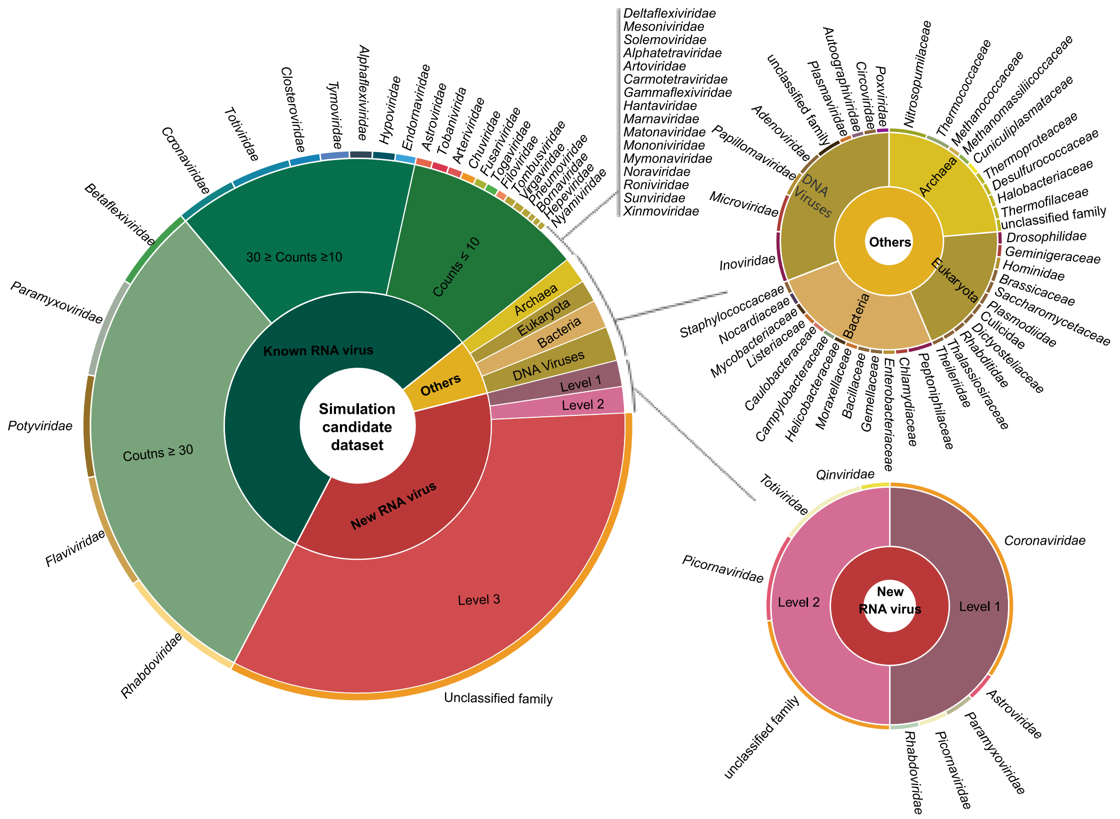


**Supplementary Figure S4. Composition of the simulation data set candidate pool.** The simulated pool includes 'known RNA viruses', 'new RNA viruses', and 'others'. The 'known RNA viruses' were derived from the NCBI RefSeq database. The 'new RNA viruses' were subdivided into three levels based on their sequence similarity to the reference database and were sourced from recent research publications. The 'others' includes DNA viruses, eukaryotes, bacteria, and archaea.

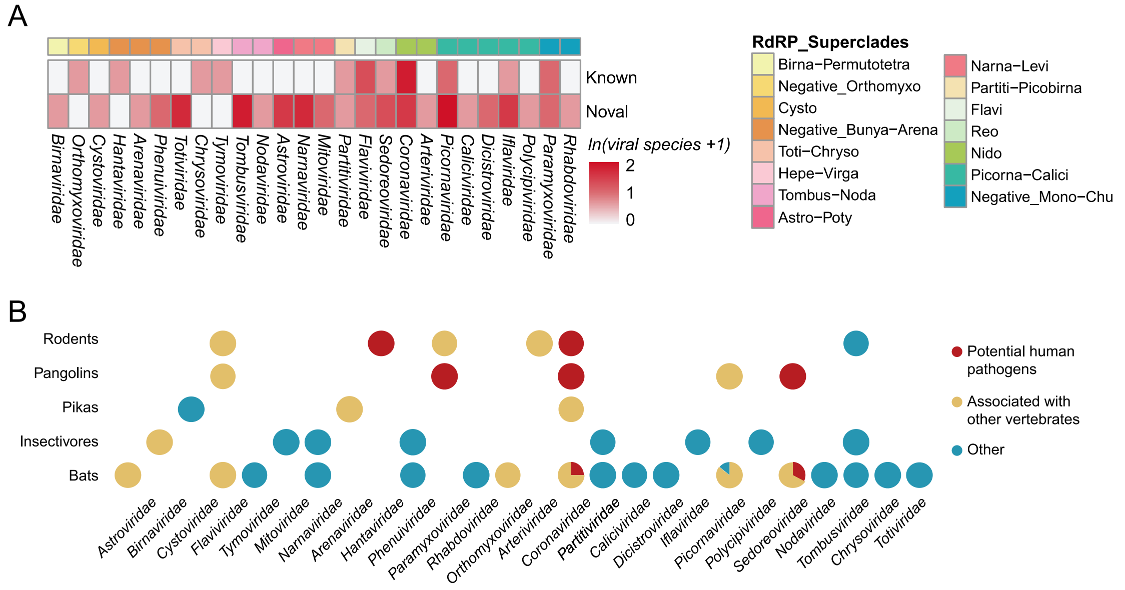


**Supplementary Figure S5. Viruses and host associations revealed by VirID. (A)** The distribution of virus species across different superclades and families identified by VirID in 20 SRA libraries. **(B)** The number of virus sequences related to humans, vertebrates, and other hosts as identified by VirID in 20 SRA libraries.

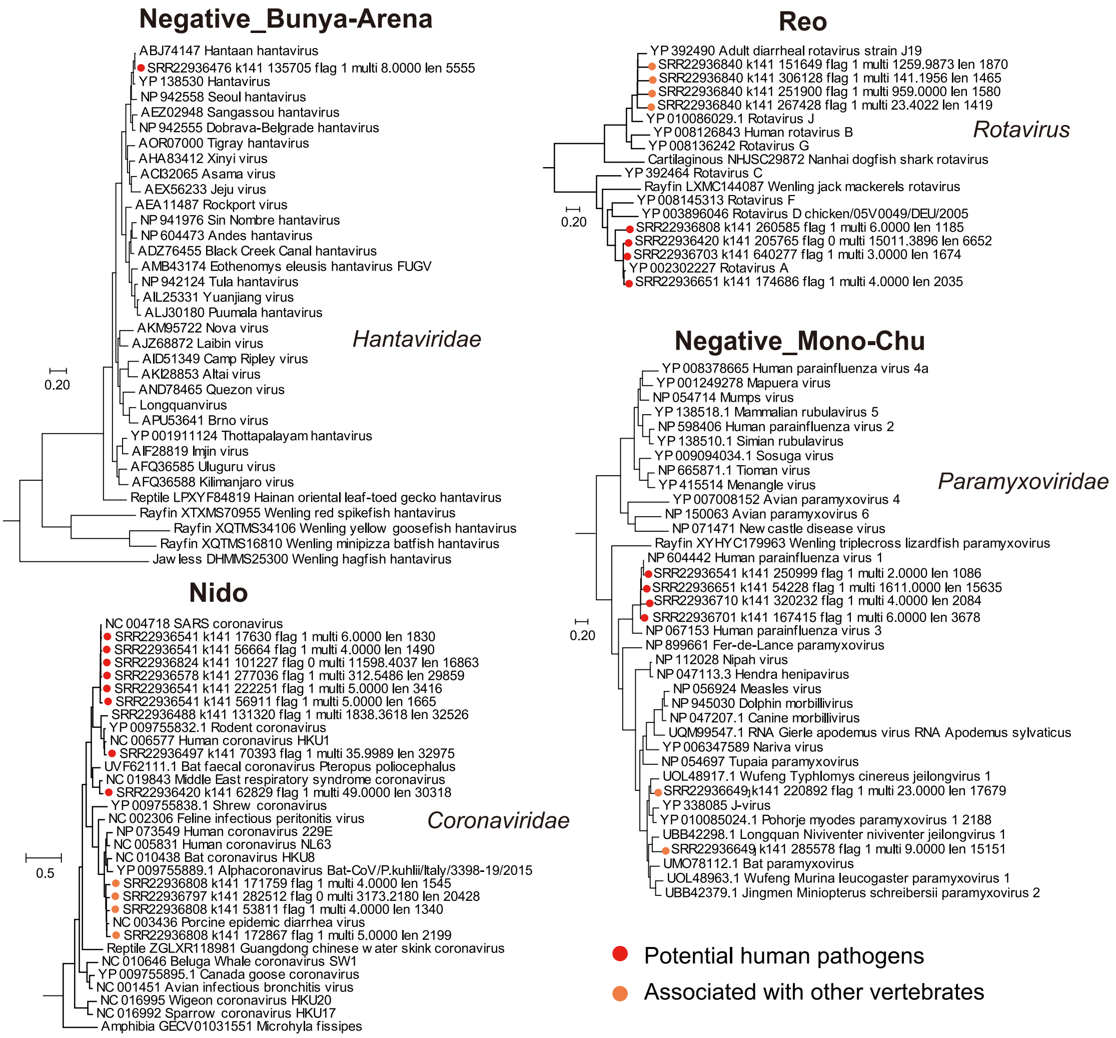


**Supplementary Figure S6. Phylogeny of viruses associated with human or vertebrates.** Phylogenetic tree shows the evolutionary relationships of the *Hantaviridae*, *Coronaviridae*, *Paramyxoviridae*, and *Sedoreoviridae* (set within their higher-level virus groups) that contain potential human pathogens. Viral sequences identified by VirID. Those labelled in red are potential human pathogens, while sequences labelled in orange are viruses associated with vertebrates.

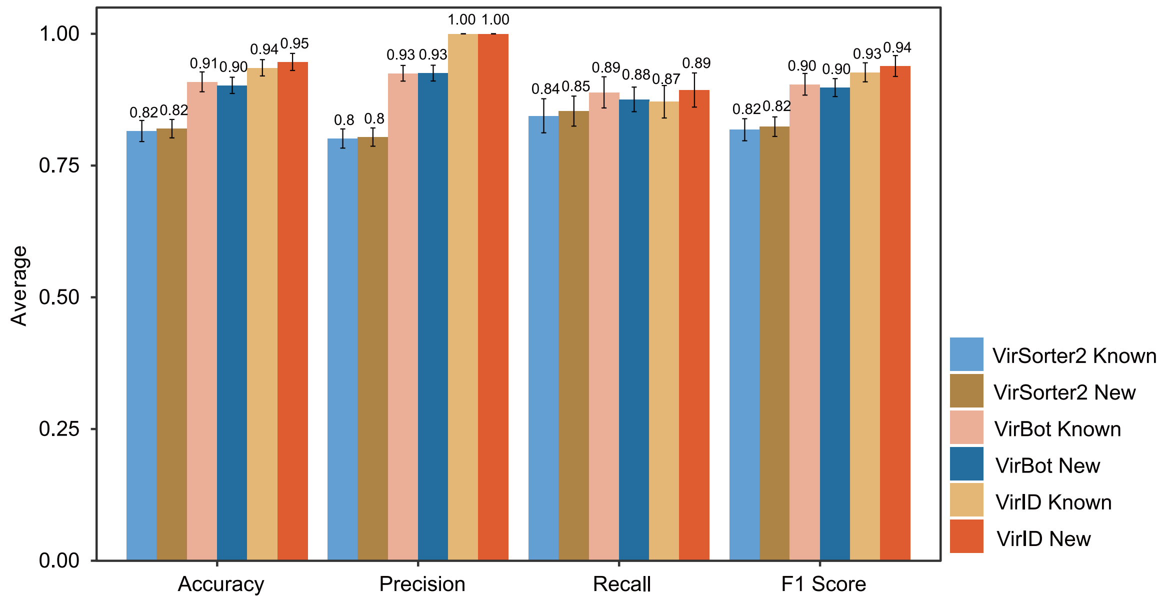


**Supplementary Figure S7. Performance of Three Tools – VirBot, VirSorter2, and VirID – in identifying 'new RNA viruses' and 'known RNA viruses'.** VirID recorded zero false positives in the identification of 'new RNA viruses'.

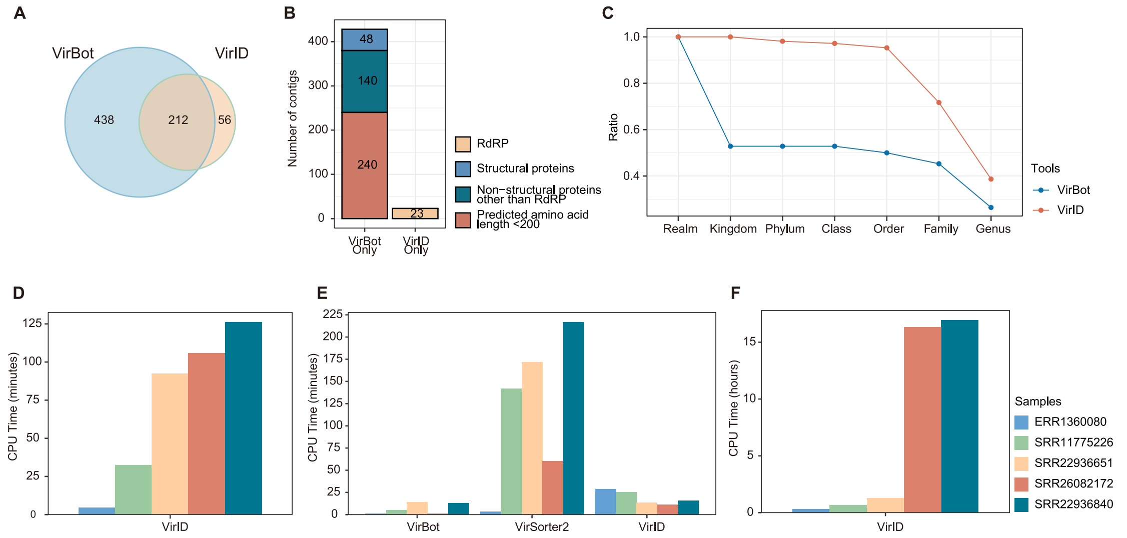


**Supplementary Figure S8. Benchmarking of different tools using real-world sequencing data. (A)** Venn diagram showing the viral contigs identified by both VirID and VirBot across 20 SRA libraries. **(B)** Bar graph depicting the annotation of unique viral genomes identified by VirID and VirBot, excluding sequences deemed false positives in VirID's phylogenetic analysis. **(C)** Line graph illustrating the percentage of sequences classified into each category. **(D)** CPU runtime estimates for the read assembly steps in VirID's analysis pipeline, based on five real-world samples with varying sequencing depths. The estimations were performed using 32 threads of the AMD EPYC 7643 CPU. Different colors represent different samples. **(E)** CPU runtime benchmark comparing VirID, VirBot, and VirSorter2 for RNA virus contig discovery across five samples, using assembled contigs as input for all three methods. The color scheme matches that of panel (D). **(F)** CPU runtime estimates for the phylogenetic inference steps in VirID's analysis pipeline. The color scheme matches that of panel (D).

### Supplementary Tables

**Supplementary Table S1. Mapping of the RdRp protein reference sequence database of RNA viruses to the ICTV taxonomic system.**

**Supplementary Table S2. Detailed information on 20 meta-transcriptomic data sets obtained from a previous study of bat, rodent, pangolin and zoo animals in China.**

**Supplementary Table S3. Detailed information on 192 public libraries from 7 studies retrieved from the SRA database.**

**Supplementary Table S4. Summary of data sets used for benchmarking based on real-world data.**

**Supplementary Table S5. Output information from the assembly and basic annotation phase.**

**Supplementary Table S6. Output information of the phylogenetic analysis section.**

**Supplementary Table S7. Detailed information on the classification results produced by VirID in seabird ticks.**

**Supplementary Table S8. Detailed information on possible host associations of viral sequences predicted by VirID in seven real-world cases.**
